# Sex-Based Transcriptomic Differences in Psoriatic Lesions: A Comprehensive Meta-Analysis

**DOI:** 10.1101/2025.05.07.652611

**Authors:** Edia Stemmer, Liat Anabel Sinberger, Tair Lax, Tamar Zahavi, Guy Shrem, Inbal Mor, Mali Salmon-Divon

**Affiliations:** Department of Molecular Biology, Ariel University, Ariel 40700, Israel; Fertility clinic, north distinct, Clalit Health Services, Israel; Maccabi Healthcare Services, Tel Aviv, Israel; Adelson School of Medicine, Ariel University, Ariel 40700, Israel

## Abstract

Psoriasis, a chronic inflammatory skin disease affecting men and women equally, presents distinct gender-based differences in severity and treatment response. While molecular mechanisms underlying psoriasis are well-studied, sex-specific differences remain largely unexplored. To address this, we conducted a meta-analysis of transcriptomic data from lesional psoriasis skin and healthy controls, comparing male and female cohorts.Our findings reveal 2,760 overlapping differentially expressed genes (DEGs) between sexes, highlighting shared pathways like IL-17 signaling and Th17 differentiation. However, sex-specific pathways emerged, including male-enriched PI3K-Akt signaling and chemokine receptor activity, and female-enriched glycolysis and AHR-NRF2 pathways. Upstream regulator analysis identified sex-specific drivers, including VEGFA activation and CFTR inhibition in males, and AHR activation and FGF21 inhibition in females. Notably, Tregs and neutrophil abundance differed by sex, aligning with disease severity trends. These results highlight critical molecular and cellular disparities, emphasizing the need for personalized, sex-specific therapeutic approaches in psoriasis management.

**Teaser:** Gender disparities in psoriatic lesions reveal transcriptomic differences and variations in neutrophil cell type abundance.

## Introduction

Psoriasis is a chronic inflammatory skin disease, affecting both men and women equally that can develop at any age(1). It occurs worldwide and imposes a significant burden on individuals and society particularly in high-income and high Socio-Demographic Index (SDI) countries in North America and Europe(2).

Psoriasis manifests in various clinical phenotypes, with chronic plaque psoriasis (psoriasis vulgaris) being the most common and easily recognizable(3). Psoriatic lesions typically appear as well-demarcated, erythematous scaly patches or thick, raised plaques with silvery scales(4). These features result from increased keratinocyte proliferation, abnormal differentiation, and incomplete cornification, leading to the retention of nuclei in the stratum corneum (parakeratosis). The lesions are also characterized by an inflammatory infiltrate of dendritic cells, macrophages, and T cells in the dermis, with neutrophils and some T cells in the epidermis, and increased vascularity with tortous capillaries contributing to the redness(5).

Despite extensive research highlighting the involvement of genetic and environmental factors—such as stress, microorganisms, drugs, trauma, and smoking—in the etiology and pathogenesis of psoriasis, the exact mechanisms remain unclear(6). However, several key immunopathologic pathways are known to be involved, including extracellular signaling pathways like TNF, IL-17, and IL-23, as well as intracellular pathways such as NF-κB, JAK-STAT and TYK2(7).

The therapeutic approach for psoriasis varies substantially, ranging from topical agents for mild cases to systemic treatments for moderate disease, and advanced biologics for severe cases targeting cytokines such as TNF-α(8).

Sex-specific manifestations of psoriasis have gained increasing attention in clinical research. While the overall prevalence is comparable between sexes, significant differences exist in disease expression and severity patterns(9). Male patients consistently demonstrate higher psoriasis area and severity index (PASI) scores(10) and require more intensive therapeutic interventions, particularly biologics(11). Recent investigations have also uncovered sex-specific variations in psoriatic arthritis presentation and progression, suggesting the need for sex-specific therapeutic approaches(12).

Sex differences in terms of hormones and gene expression are considered to be the main reasons for differences in diseases(13, 14). Additionally, variability in drug metabolism along with social and cultural factors related to gender differences, may also have a major impact on the development and experience of psoriasis.

Recognizing the growing importance of biological sex and gender differences in disease phenotype and drug response, we conducted a mega-analysis of transcriptomic data derived from lesional biopsies of psoriasis patients and healthy controls, including both male and female cohorts. To our knowledge, this is the first meta-analysis to investigate the molecular transcriptomic differences between males and females in psoriasis.

## Results

### Study data and Sample Distribution

Following the PRISMA recommendations in the GEO and ArrayExpress databases, our search identified 13 studies containing transcriptomic data of 342 skin samples that met our inclusion criteria (Figure 1). Only four studies provided detailed information regarding sex. For the remaining nine studies (marked with an asterisk in Table 1), we used a logistic regression model based on gene expression from specific markers to infer sample sex. MDS plots before and after the inference are depicted in Supplementary Figure S2. Table 1 provides a detailed summary of the distribution of psoriasis and control samples by sex for each study.

**Table 1:** Study datasets by disease status and sex following sex determination.

| No. | GEO\ArrayExpress Study | Psoriasis |  | Control |  |
| --- | --- | --- | --- | --- | --- |
|  |  | Female | Male | Female | Male |
| 1 | EMTAB6556 | 2 | 8 | - | - |
| 2 | GSE117405* | 8 | 11 | 8 | 1 |
| 3 | GSE121212* | 13 | 13 | 19 | 15 |
| 4 | GSE171012* | 6 | 8 | 6 | 4 |
| 5 | GSE176279* | - | 2 | - | - |
| 6 | GSE183820 | - | 1 | - | - |
| 7 | GSE205748* | - | - | 5 | 4 |
| 8 | GSE249936 | 3 | 5 | - | - |
| 9 | GSE41745* | 2 | 1 | - | - |
| 10 | GSE47944 | 3 | 5 | 5 | - |
| 11 | GSE54456* | 25 | 60 | 31 | 49 |
| 12 | GSE67785 | 6 | 8 | - | - |
| 13 | GSE83645* | 1 | 4 | - | - |
| **Total = 342** |  | **69** | **126** | **74** | **73** |

\* Studies had sex inferred via RNA-seq using a logistic regression model and six sex-specific genes.

**Figure 1:**
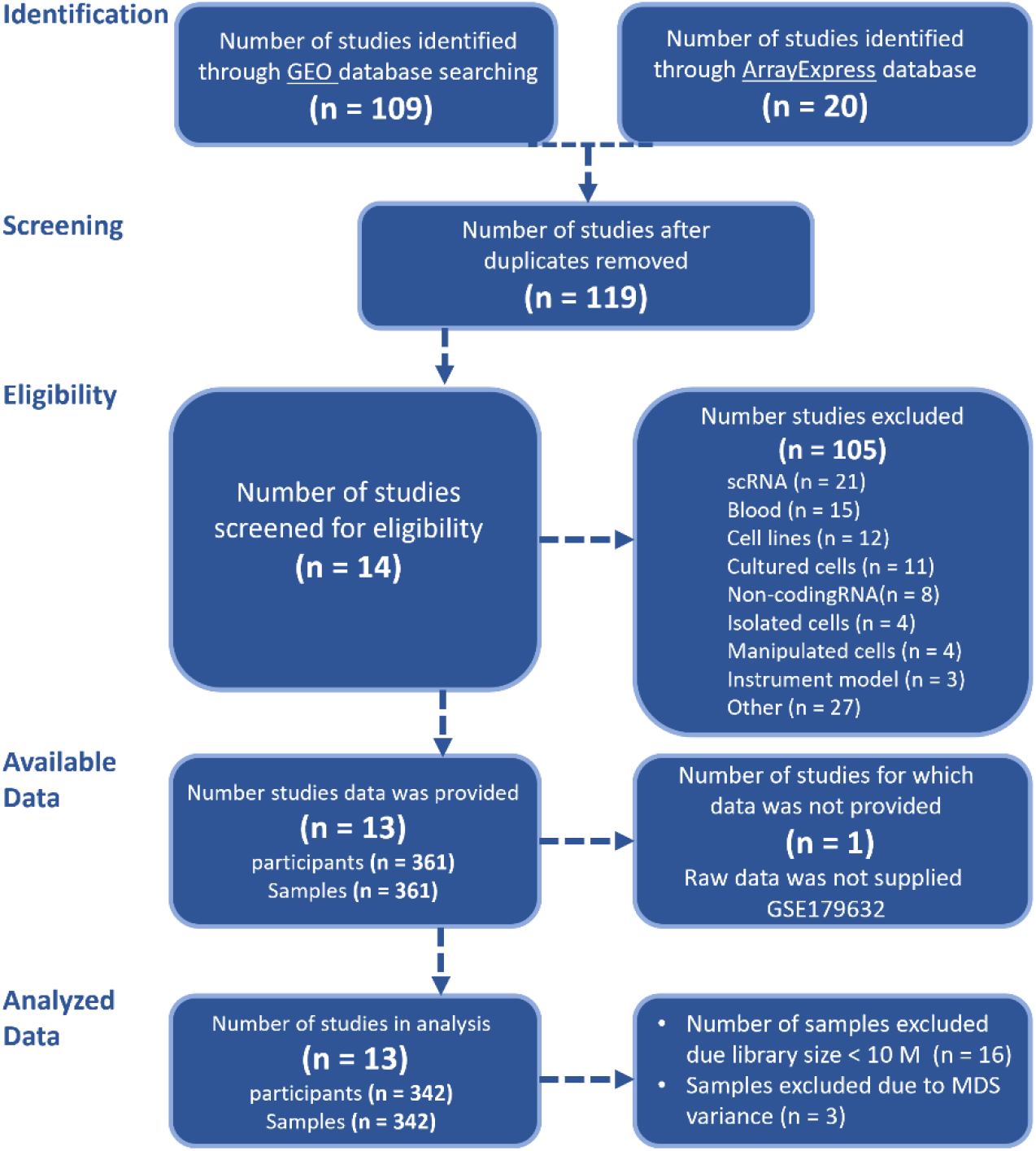
PRISMA flow diagram. PRISMA: Preferred Reporting Items for Systematic Reviews and Meta-Analyses

### MDS Analysis Reveals Disease and Sex-Driven Expression Variation

MDS analysis of biopsy expression levels was conducted after the removal of study batch effect, revealing that the primary factor driving variation was the disease status, distinguishing between lesional psoriasis biopsies and control skin biopsies (Figure 2A). Subsequently, within each group, the next factor influencing variation was the sex status of the biopsies (figure 2B).

**Figure 2:**
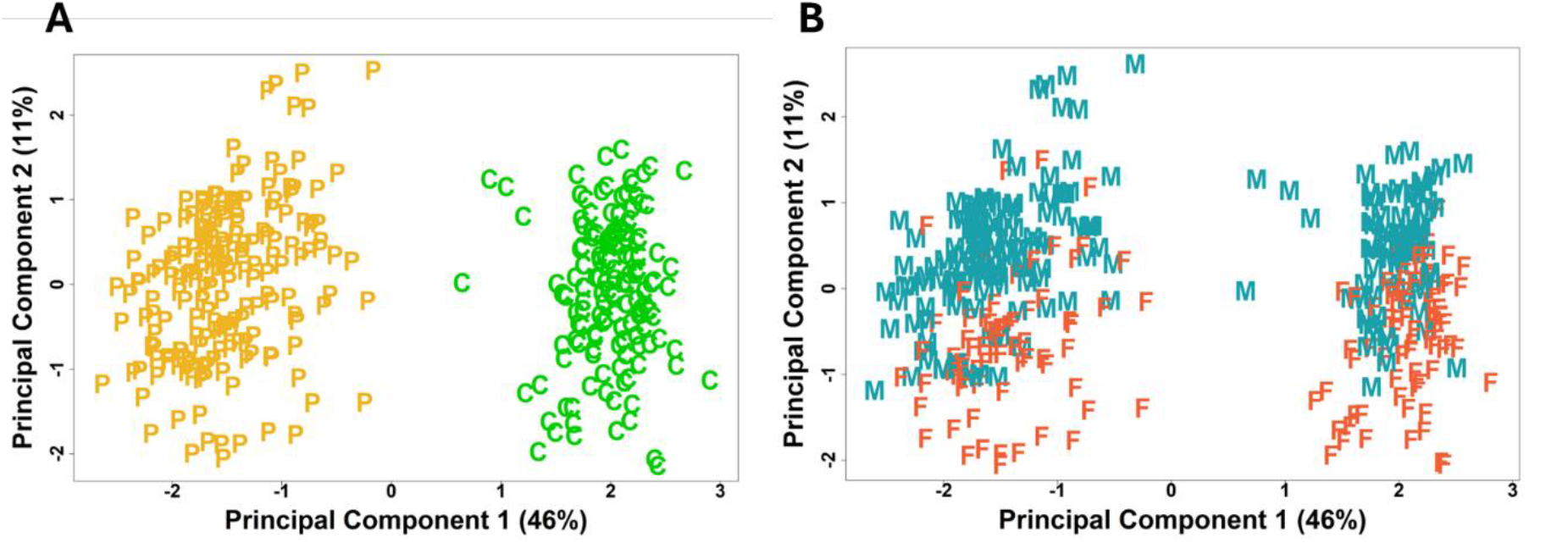
MDS analysis. MDS analysis of normalized biopsy expression data, with batch effects removed. (A) Disease status—Psoriasis (P, yellow) vs. Control (C, green), and (B) Sex status—Female (F, orange) vs. Male (M, turquoise).

### Pathway Enrichment and Upstream Regulator Analysis of DEGs

Differential expression gene (DEG) analysis revealed 3,309 DEGs (Full list of DEGs is available upon request) in females with psoriasis compared to female controls, and 3,221 DEGs in males with psoriasis compared to male controls. Of these, 2,760 DEGs were common to both comparisons, with consistent directionality (upregulated or downregulated in both). Notably, the S100A family and IL17 genes were upregulated in both male and female psoriasis groups. Additionally, 549 DEGs were unique to females, while 461 DEGs were unique to males. The directional distribution of DEGs is illustrated in Figure 3A-C, showing the Venn diagrams sections of the two comparisons. Metascape analysis was conducted for each section of the Venn diagrams, examining pathways associated with the overlapping DEGs in male and female samples and those unique to each sex-specific comparison (Psoriasis male vs. Control male and psoriasis female vs. Control female). For the overlapping DEGs, key pathways identified included the “Th17 cell differentiation pathway”, “IL-17 signaling pathway”, “Cytokine-cytokine receptor interaction”, as well as pathways related to calcium regulation such as the “Calcium signaling pathway” and “Vitamin D receptor pathway” (Figure 3A). DEGs unique to psoriasis females were associated with sugar metabolism pathways, including “aerobic glycolysis” and “fructose and mannose metabolism”, as well as oxidative stress pathways like “NRF2” and “FOXA2”. “NF-kappa B signaling”, crucial for inflammation and immune activation, was also implicated. (Figure 3B). For the male psoriasis comparison, distinct DEGs were found in pathways like “Cell adhesion molecules”, “Cytokine-cytokine receptor interaction”, “Tight junction” and “PI3K-Akt signaling pathway” emphasizing the roles of inflammation, barrier function disruption, and cell proliferation in driving the pathology of psoriasis in males. (Figure 3c).

**Figure 3:**
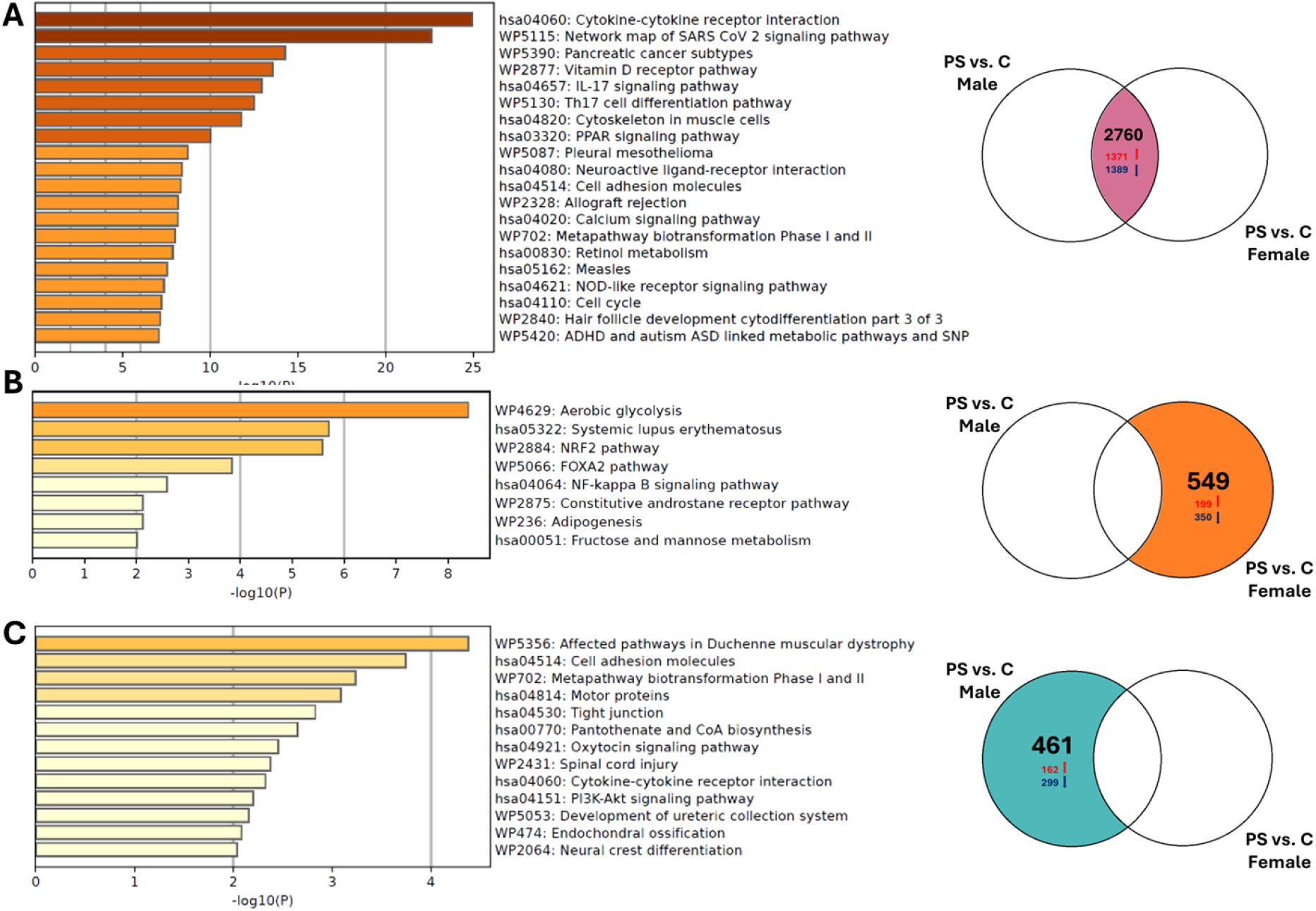
Metascape and Venn diagram of psoriasis DEGs. Metascape analysis is provided for each section of the Venn diagram, detailing the enrichment of biological terms and pathways specific to the gene sets. The Venn diagram shows DEGs between psoriasis (PS) and control (C) for both males and females, including overlaps. Arrows indicate DEG direction: red for upregulated and blue for downregulated. (A) Pathways for overlapping DEGs. (B) Pathways for DEGs unique to female psoriasis (not in males). (C) Pathways for DEGs unique to male psoriasis (not in females).

IPA upstream regulator (UR) analysis, using an absolute Z score of 2 or greater and a p-value less than 0.05, identified regulators across various molecular types. Table 2 highlights regulators of molecule types related to immune responses (cytokines, transmembrane receptors), proliferation and differentiation (growth factors), and other metabolic and intracellular signaling pathways relevant to psoriasis pathogenesis that were found exclusively in one sex. It is worth noting that for all the regulators that were common to both sexes, the directionality of the effect was the same in both groups; if a regulator was activated in males, it was also activated in females, and the same applied to inhibited regulators.

**Table 2:** Upstream regulators unique to each sex in psoriasis vs. control. Red denotes activated UR, blue denotes inhibited UR.

| Sex | Transcription regulator | Ligand-dependent nuclear receptor | Growth factor | Transmembrane receptor | Cytokine | Transporter | Ion channel |
| --- | --- | --- | --- | --- | --- | --- | --- |
| Male | TBX21,STAT5A,REL,CREB1,KLF5,ACTN4,TOX,YAP1,YBX1,ATF6,C1QBPTBX2,STAT4,ETV4,MITF,ETS2,ELK1,VAV2,PRDM1,ATF3,RBL1,ARID1A,GATA6TRPS1,PPARGC1A,SATB1,MXD1,MYOCD | PGR,NR3C1 | FGF2,VEGFA,NRG1,TGFB2 | TNFRSF1B, Klrk1,IL2RG,IL11RA,ITGAV,ICAM1,CD80,PDCD1,LILRB1 | TNFSF15,TNFSF14,CCL3,CXCL10,IFNE,CNTF | CEACAM1,TAP1,SLC6A4 | CFTR |
| Female | STAT3,SP1,MSC,FOSL1,SOX2, POU2AF1,SOX11,GLIS1,COMMD3-BMI1,PAX1 | AHR | FGF21 | TNFRSF1A,TNFRSF10A,CHRNA1,LEPR |  | NPC1 | KCNJ10 |

### Sex-Specific DEGs

Ten DEGs were identified in the interaction analysis between male and female psoriasis patients, as shown in Table 3. Among these, SYT6, PSG4, NELL2, GPR15, MAP1LC3C and C10ORF67 are protein-coding genes, CYP4Z2P, RN7SL5P and CLCA3P are pseudogenes, and LINC01698 is a long non-coding RNA (lncRNA). None of these genes are located on the sex chromosomes. Table 3 provides detailed information about these genes, obtained from the GeneCards database (www.genecards.org)(15), demonstrating a high representation of proteins that are relevant to calcium signaling or intracellular signaling. The expression of these interaction genes is presented in boxplots in Supplementary Figure S3. GSEA of genes ranked based on their interaction fold change, identified similar pathways that were detected as differentially enriched in psoriasis males as mentioned above. For example, bioenergetic pathways such as “Cell Adhesion Molecules,” “Glycolysis/Gluconeogenesis,” and “Oxidative Phosphorylation” were less enriched in male psoriasis, while proinflammatory pathways like “IL-17 Signaling” and “Chemokine Receptors Bind Chemokines” were more enriched. These pathway analyses are illustrated in Supplementary Figure S4.

**Table 3:** Sex-specific DEGs in Psoriasis. Expression Status indicates changes in gene expression direction, based on comparisons between psoriasis and control samples, specific to each sex: statistically downregulated or upregulated in male (M) or in female (F) samples, or in both (F, M).

| Gene | Cytogenetic location | Expression status found in our analysis | Description and role | Related pathways |
| --- | --- | --- | --- | --- |
| SYT6 | 1p13.2 | Downregulated (M) | (Synaptotagmin 6): includes a transmembrane domain and a cytoplasmic region composed of 2 C2 domains and are involved in calcium-dependent exocytosis of synaptic vesicles. | calcium ion binding<br>identical protein binding |
| NELL2 | 12q12 | Upregulated (F) | (Neural EGFL Like 2): Glycoprotein, Ca<sup>2+</sup>-binding sites, protection of cells from cell death-inducing environments. prevented ER stress-mediated apoptosis | Nervous system development<br>calcium ion binding |
| PSG4 | 19q13.31 | Upregulated (M)<br>Downregulated (F) | (Pregnancy Specific Beta-1-Glycoprotein 4): May play a role in regulation of the innate immune system | Response to elevated platelet cytosolic Ca<sup>2+</sup><br>Cell surface interactions at the vascular wall |
| CLCA3P | 1p22.3 | Upregulated (M, F) | (Chloride Channel, Calcium Activated, Family Member 3 Pseudogene): Part of the calcium-sensitive chloride conductance protein family. | *Pseudogene* |
| GPR15 | 3q11.2 | Upregulated (M, F) | (G Protein-Coupled Receptor 15): Acts as a chemokine receptor | GPCR downstream signaling<br>Class A/1 (Rhodopsin-like receptors)<br>transmembrane signaling receptor activity |
| MAP1LC3C | 1q43 | Downregulated (M, F) | (Microtubule Associated Protein 1 Light Chain 3 Gamma): Involved in autophagy that plays an important role in cellular maintenance and development | Selective autophagy<br>microtubule binding |
| CYP4Z2P | 1p33 | Upregulated (M, F) | (Cytochrome P450 Family 4 Subfamily Z Member 2 Pseudogene): Including heme binding activity; iron ion binding activity; and monooxygenase activity | *Pseudogene* |
| RN7SL5P | 9p23 | Upregulated (F) | (RNA, 7SL, Cytoplasmic 5 Pseudogene): Non-coding RNA. | *Pseudogene* |
| LINC01698 | 1q32.2 | Downregulated (M, F) | (Long Intergenic Non-Protein Coding RNA 1698): Long non-coding RNA with regulatory functions. | *lncRNA* |
| C10orf67 | 10p12.2 | Downregulated (M, F) | Chromosome 10 Open Reading Frame 67 | *No data available* |

### Differential Cell Type Enrichment Between Sexes in Psoriasis

To explore possible sex-based variation in cell populations within the samples we used the xCell webtool to perform cell type enrichment analysis on bulk gene expression data, normalized for gene length, across four groups: psoriasis males, psoriasis females, control males, and control females. The heat map displayed in Figure 4A shows the average scores for each cell type across these groups. Psoriasis was associated with significant changes in various cell populations compared to control with increased immune and microenvironment scores and decreased stromal scores. While the overall patterns in the heatmap appear similar between sexes in both the psoriasis and in control groups, a more detailed analysis reveals significant sex-based variations within the psoriasis group, as illustrated in Figure 4B.

**Figure 4:**
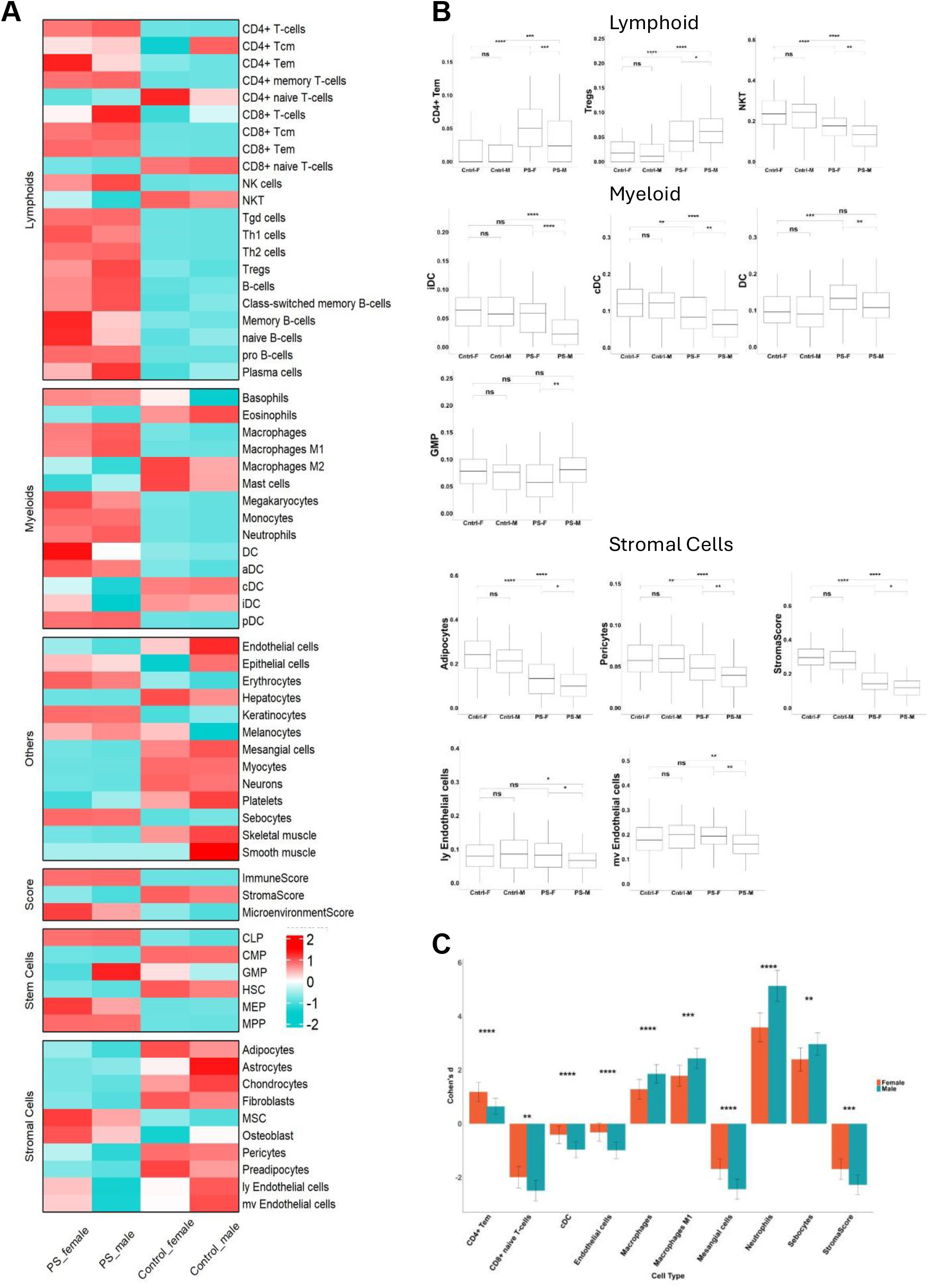
Cell type enrichment analysis. (A) Heatmap of cell type enrichment analysis showing the average xCell scores for each group: PS-Female, PS-Male, Control-Female, and Control-Male. (b) Boxplots illustrating significant sex-specific differences in psoriasis. Groups compared include PS-Female (PS-F), PS-Male (PS-M), Control-Female (Cntrl-F), and Control-Male (Cntrl-M). Statistical significance was assessed using Dunn’s test, with p-values adjusted using FDR correction. Asterisks indicate significance levels. (C) Barplots displaying Cohen’s D scores for cell types, representing the effect size of differences between groups. Cohen’s D was calculated separately for females (PS-F vs. Cntrl-F) and males (PS-M vs. Cntrl-M). For each comparison, the variances of Cohen’s D were estimated, and a z-statistic was computed to evaluate the differences between male and female effect sizes. P-values were derived from the z-statistics and corrected using FDR. The y-axis shows Cohen’s D values, while the x-axis lists cell types. Error bars represent confidence intervals. Significance levels: *p < 0.05, **p < 0.01, ***p < 0.001, ****p < 0.0001, and “ns” for non-significant.

Regulatory T cells (Tregs) were significantly higher in male compared to female, while NKT cells and CD4+ Tem cells were lower in males. Among myeloid cells, conventional dendritic cells (cDC), immature dendritic cells (iDC), and overall dendritic cells (DC) were reduced in male, whereas granulocyte-monocyte progenitors (GMP) were increased. Regarding stromal cells, most are significantly reduced in male psoriasis compared to females (Figure 4B).

Next, we sought to determine whether there are significant sex-based differences in microenvironment responses to psoriasis and how large those differences are. For this aim, we used Cohen’s D calculation, assessing how much the xCell enrichment scores differ between the psoriasis and control groups for each sex. The results of this analysis are shown in Figure 4C.

We observed elevated scores for neutrophils and M1 macrophages in male psoriasis, alongside reduced scores for cDCs. Mesangial cells significantly decreased in males, while sebocytes increased. CD4+ Tem cells were the only cell type that significantly increased in female psoriasis.

These results highlight the complex and sex-specific cellular alterations in psoriasis, underscoring the importance of considering sex when analyzing immune and stromal cell populations in this disease.

## Discussion

Psoriasis is a chronic inflammatory skin disease affecting both men and women. Although many studies have examined the molecular mechanisms involved in psoriasis by transcriptomic analyses of healthy and psoriatic skin, sex-based differences remain largely unexplored, despite evidence suggesting that males often experience more severe disease(9). To uncover the role sex plays in psoriasis, we compared gene expression between lesional psoriatic skin and healthy control skin from both males and females, using uniformly reprocessed samples from 13 different studies for a comprehensive analysis.

MDS analysis showed sex was second only to disease status as the main driver of gene expression differences, emphasizing its role in shaping disease outcomes and treatment responses.

The differential gene expression analysis in psoriasis highlights significant similarities and sex-specific differences in the underlying mechanisms of the disease. The identification of 2,760 overlapping DEGs between male and female psoriasis patients suggests the involvement of a shared set of genes in the pathogenesis of psoriasis, pointing to a common core pathways. These DEGs are notably linked to psoriasis-related pathways, such as Th17 cell differentiation, IL-17 signaling, and cytokine-cytokine receptor interactions, which are central to the inflammatory processes driving psoriasis. Furthermore, the involvement of vitamin D receptor and calcium signaling pathways also aligns with the importance of calcium metabolism in maintaining keratinocyte function(16, 17). Disrupted calcium gradients in the epidermis, often seen in psoriasis, are critical in impairing keratinocyte differentiation and skin barrier function(18). Moreover, the enriched pathway “Chemokine receptors bind chemokines” identified in the male group is associated with intracellular calcium fluxes, which regulate chemotaxis and inflammatory responses(19). Other pathways identified specifically in the male group related to tight junctions and cell adhesion molecules suggesting alterations in keratinocyte barrier integrity, which may also be tied to abnormal calcium regulation. In contrast, female-specific DEGs were enriched for bioenergetics pathways, including glycolysis and fructose metabolism, which are linked to calcium transients and have been observed to enhanced osteoblasts proliferation(20). This enrichment may indicate a metabolic adaptation in female psoriasis. Indeed increased expression of glycolysis-related genes is associated with disease severity modulation(21).

In the context of metabolic changes, we found the PI3K-Akt signaling pathway to be elevated in the male group. This pathway is activated in the keratinocytes of psoriatic lesions and higher expression of PI3K may cause Akt hyperactivity, further promoting the proliferation of keratinocytes(7). In osteoblasts, calcium transients triggered by high extracellular calcium levels, have been shown to activate glycolysis via AKT-related signaling promoting cell proliferation (20). Given the known link between glycolysis and psoriasis, and the observation that female psoriasis is associated with glycolysis-related pathways, we suggest that sex-specific factors, including calcium regulation, may influence the pathogenesis of psoriasis through these metabolic pathways.

The upstream regulator (UR) analysis revealed regulators specific to either males or females. A recent study on ion channels in psoriasis pathophysiology emphasizes CFTR’s role in epidermal cells, particularly in regulating fluid balance, reducing inflammation, and promoting keratinocyte differentiation(22, 23). In our study, CFTR inhibition was observed in male psoriasis, supporting a potentially more severe disease course in males. Activated transcription factors in males included TBX21 and STAT5A, which drive Th1 immune responses and cellular proliferation(24, 25). Growth factors such as VEGFA were also upregulated in our analysis. VEGFA is highly expressed in psoriasis lesions and linked to disease severity(26–28). The activation of VEGFA in male psoriasis could suggest a more severe pathology, aligning with male-specific pathways of inflammation and immune cell recruitment. In contrast, female-specific regulators included the activation of potassium ion channel KCNJ10, which plays a role in epithelial repair, particularly in cell migration and proliferation(29). Furthermore, female psoriasis is marked by the activation of ligand-dependent nuclear receptors like AHR along with the inhibition of growth factors such as FGF21, which are involved in metabolic regulation and oxidative stress responses. The NRF2 pathway was enriched in our female-specific analysis. The AHR-NRF2 axis is likely anti-inflammatory, reducing the production of proinflammatory cytokines. AHR agonists, such as tapinarof, are currently used as therapeutic agents in psoriasis(30). FGF21, a key regulator of keratinocyte migration and differentiation during wound healing(31), is typically elevated in the serum of psoriasis patients, correlating with disease severity(32). However, in our data, FGF21 was inhibited in female psoriasis, potentially indicating less severe disease. In summary sex-specific upstream regulators highlight distinct molecular drivers of psoriasis in males and females, suggesting that therapeutic approaches should be tailored to these differences.

We identified 10 genes that showed an interaction between sex and disease, indicating that psoriasis influences these genes differently in males and females. Of these, six are known to be associated with psoriasis. SYT6, located in the PTPN22 region, is linked to early-onset psoriasis(33) and is differentially methylated in psoriatic lesions(34). NELL2, a calcium binding glycoprotein, is deregulated in psoriatic substitutes and found to be increased in atopic dermatitis (AD) compared to psoriasis. Upregulated by Th2 cytokines(35), NELL2 prevents ER stress-induced cell death and promotes cell survival(36) by activating the ERK1/2 pathway, which is overactive in psoriasis lesions(37). This activation supports cell survival and differentiation(7, 38). Notably, NELL2 is upregulated in female psoriasis, suggesting a female-specific pathogenesis. PSG4 was found to be highly expressed in male psoriasis in our data and is associated with generalized pustular psoriasis (GPP), a more severe form of psoriasis characterized by neutrophil/monocyte dominance and extensive systemic manifestations(39). GPR15 and its ligand, GPR15LG, are involved in skin lymphocyte homing and keratinocyte proliferation, with GPR15LG significantly elevated in psoriatic lesions(40–42). MAP1LC3C showed potential links to autophagy, being downregulated in psoriatic lesions compared to normal skin(43). CYP4Z2P, a pseudogene, was upregulated in psoriasis lesions and linked to inflammatory skin diseases(44). Additionally, several other pseudogenes and noncoding genes, including RN7SL5P, CLCA3P, and LINC01698, were identified, although their functions are not well understood. However, CLCA3P is part of the CLCA protein family, which regulates chloride channels and plays an important role in keratinocyte functions such as proliferation and apoptosis(45). Finally, C10orf67, associated with sarcoidosis—an inflammatory condition affecting the skin through granuloma formation—was also identified(46). These findings highlight distinct molecular mechanisms that may contribute to sex-specific differences in psoriasis pathogenesis.

Our analysis revealed Tregs, were elevated in psoriasis lesions, with a more pronounced increase in males. Tregs are known for their anti-inflammatory properties and critical role in maintaining immune homeostasis to prevent autoimmune diseases(47). The overall rise in Tregs we identified in psoriatic lesions is consistent with previous studies and it is hypothesized that their function may be impaired in psoriasis, leading to an increase in Th17 cells and contributing to disease pathogenesis. A correlation between Treg levels and disease severity has also been suggested, though the exact role of impaired Treg function in psoriasis remains unclear(48). A recent study highlighted differences in expression of Treg markers between psoriasis and healthy samples(49). We propose that Tregs may also exhibit distinct expression markers between males and females, potentially affecting disease pathogenesis.

Several studies have explored the role of DC subtypes in the development of psoriasis. cDCs can be classified into two types: CD1c+ DCs and CD141+ DCs. CD1c+ DCs are reduced in both non-lesional and lesional skin in psoriasis, while CD141+ DCs are increased in these areas(50, 51). In our analysis of lesional skin, we observed a greater reduction of cDCs in male psoriasis patients. Additionally, while iDCs are typically increased in psoriasis lesions, our data indicate a lower abundance of iDCs in males. These differences may indicate a sex-dependent effect on DC function.

Stromal cells, including fibroblasts and pericytes, were reduced in male psoriasis, indicating a potentially more severe disease progression due to impaired tissue repair and disrupted skin integrity. Neutrophils play a crucial role in psoriasis, serving as a histopathologic hallmark in lesions and contributing to disease development through processes such as respiratory burst, degranulation, and the formation of neutrophil extracellular traps (NETs)(52). Cohen’s D analysis revealed a greater increase in neutrophils in male psoriasis patients compared to females. The abundance of neutrophils is particularly characteristic of severe forms such as generalized pustular psoriasis, and neutrophil depletion has shown efficacy in alleviating symptoms in pustular psoriasis in patients that did not respond well to conventional treatments(53). The neutrophils processes like degranulation and NETs formation are triggered via different stimuli, including cytosol calcium levels and ROS production which can be influenced by sex hormone(54). Altogether our finding that psoriatic lesions in males have greater increase in neutrophils may reflect the increased severity of the disease in male patients.

This study has several limitations. Not all studies included in the meta-analysis provided data on age, race, disease duration, medication, lifestyle factors, or the location of biopsies. Additionally, PASI score was missing in most cases. These factors may significantly influence gene expression profiles and limit the generalizability of the findings. Nevertheless, all studies collected samples from lesional areas, allowing our findings to focus on affected regions and provide insights into key aspects of psoriasis pathogenesis. Another limitation concerns the control biopsies, for which information on anatomical location and donor history was unavailable. This lack of data may have impacted the results. Despite these limitations, the clear separation between control and lesional samples observed in MDS analyses supports the validity of the distinction between the two groups.

In conclusion, our study underscores the importance of sex differences in psoriasis pathogenesis. By identifying sex-specific DEGs, pathways, upstream regulators, and immune cell types, we provide insight into the distinct molecular and cellular features of psoriasis in males and females. These findings could guide the development of more targeted therapeutic strategies, potentially improving treatment outcomes for both sexes. Future research should further explore these sex-specific characteristics and their clinical relevance in larger, more diverse cohorts.

## Materials and Methods

### Search methods and Selection criteria

A comprehensive search of the Gene Expression Omnibus (GEO)(55) and ArrayEpress (AE)(56) databases was conducted on June 2, 2024. The search terms included: ((“psoriasis”) OR (“psoriatic”)) AND (“high throughput sequencing"[Platform Technology Type]) AND “homo sapiens"[Organism] AND (“RNA-seq” OR “RNAseq” OR “RNA sequencing”). This systematic review was carried out in accordance with the PRISMA (Preferred Reporting Items for Systematic Reviews and Meta-Analyses) guidelines. The flowchart illustrating the study selection process is shown in Figure 1. Studies were included in the mega-analysis only if they met the following inclusion criteria: availability of RNA-seq raw data, transcriptomes derived from whole lesional or healthy control skin tissue samples (punch biopsy), sequencing conducted without additional manipulations (e.g., tissue culture, isolated cells), use of the Illumina sequencing platform, and availability of corresponding metadata. Studies utilizing microarray, single-cell RNA-seq (scRNA-seq), or small RNA-seq platforms were excluded.

Notably, control samples were obtained from normal, healthy individuals, including samples collected following cosmetic surgery, ensuring they represented non-psoriatic skin tissue.

### RNA Sequencing Data Collection and Processing

RNA sequencing data from the selected studies were uniformly reprocessed. Raw data were downloaded using the SRA-Toolkit (https://hpc.nih.gov/apps/sratoolkit.html). In brief, the fasterq-dump tool was used to download FASTQ files, followed by adapter trimming and quality checks with Trim Galore (www.bioinformatics.babraham.ac.uk/projects/trim_galore/) and FASTQC (https://www.bioinformatics.babraham.ac.uk/projects/fastqc/), respectively. High-quality reads were aligned to the reference genome (GRCh38) using HISAT2(57), and the number of reads mapped to each annotated gene was counted with featureCounts(58). Only samples with a library size greater than 10 million counts were retained for statistical analysis, and genes with low read counts were filtered out using the ‘filterByExpr’ function from the edgeR package(59).

### RNA-seq Sex Inference

To determine the biological sex of samples where the sex was unknown, we cloned the SexInference GitHub repository (https://github.com/SexChrLab/SexInference.git) to develop and test a logistic regression model with Lasso regularization for inferring biological sex from RNA-seq data. The model incorporated the expression levels of six sex-specific genes: XIST (female-specific) and five Y-linked genes (EIF1AY, KDM5D, UTY, DDX3Y, and RPS4Y1). We trained and tested the model using gene expression data from the GTEx project, which includes known biological sex information. The logistic regression model was optimized using cross-validation with cv.glmnet to select the best regularization parameter (lambda). A threshold of 0.5 was used for classifying biological sex based on the model’s predicted probabilities, with predictions above indicating male and below indicating female. The trained model was then applied to our RNA-seq experimental data, where the true sex was not known, to infer the biological sex of the samples.

### Differential Expression Analysis

Normalization and differential gene expression analysis were conducted using the edgeR(60) and limma(61) R packages. Briefly, raw read counts were normalized using the TMM method, followed by a voom transformation to approximate a normal distribution. This transformation produced a new dataset with values in logCPM (log2 counts per million), which were then used with classic normal-based statistical methods and workflows to detect differentially expressed genes (DEGs). Prior to initiating the differential expression analysis, we performed Multidimensional Scaling (MDS) analysis to visualize batch effects associated with the dataset in a reduced-dimensional space. We found that the “study source” factor had the largest contribution to the variation (Supplementary Figure S1). Consequently, the study feature was included as a covariate in the model to account for its effect. The dataset was divided into four groups: psoriasis male (PS_male), psoriasis female (PS_female), healthy control male (Control_male), and healthy control female (Control_female). DEG comparisons were performed separately for Psoriasis vs. Control male and Psoriasis vs. Control female. Additionally, we conducted a DEG analysis to study the interaction effect between sex and disease, represented by the contrast matrix interaction = (PS_male – Control_male) – (PS_female – Control_female). Statistically significant DEGs were defined as those with |log2 fold change (FC)| > 1 (either upregulated or downregulated) and an adjusted P value (Benjamini-Hochberg) < 0.05.

### Pathways enrichment analysis using Metascape, IPA and WebGstalt

We performed pathway enrichment analysis using Metascape, IPA, and WebGestalt to explore biological processes and pathways relevant to psoriasis. First, we created Venn diagrams (https://bioinformatics.psb.ugent.be/webtools/Venn/) to compare the DEGs obtained from the two comparisons: control vs. psoriasis female and control vs. psoriasis male. This analysis allowed us to identify overlapping DEGs, as well as those unique to each comparison. We then used Metascape(62) (http://metascape.org/gp/index.html#/main/step1), a web-based tool integrating data from sources like Reactome, Canonical Pathways, BioCarta, GO Biological Processes, Hallmark Gene Sets, and KEGG Pathways, to perform enrichment analysis on the DEGs from each section of the Venn diagram. Metascape provided interactive visualizations and summary reports to highlight overrepresented biological terms and pathways. We displayed -log10(p-value) values greater than 2, corresponding to p-values less than 0.01, which are considered statistically significant. Additionally, we used Ingenuity Pathway Analysis (IPA)(63) (QIAGEN Inc., https://www.qiagenbioinformatics.com) to analyze upstream regulators associated with the DEGs from each comparison (males or females). IPA identified significant upstream regulators based on their ability to influence the expression of the DEGs. Negative z-scores indicated inhibition of regulators, while positive z-scores indicated activation. Regulators with p-values < 0.05 and absolute z-scores ≥ 2 were considered statistically significant. Finally, Gene Set Enrichment Analysis (GSEA) was conducted using WebGestalt (https://www.webgestalt.org, PMID: 38808672) to identify pathways enriched in gender-specific psoriasis interactions. Genes were ranked based on differential expression analysis of the interaction term (PS_male – Control_male) – (PS_female – Control_female), highlighting pathways enriched or depleted in psoriasis males compared to females. Predefined gene sets were analyzed, and significant pathways were identified using FDR correction (adjusted p < 0.05). Results were visualized through barplots and enrichment plots.

### Cell type enrichment analysis using xCell

xCell(64) (https://xcell.ucsf.edu/) is a web tool utilized for cell type enrichment analysis, providing enrichment scores (Ess) for 64 immune and stromal cell types based on gene expression data. To perform the analysis, we first normalized the gene expression data using the Reads Per Kilobase per Million mapped reads (RPKM) method, which adjusts for gene length. We then addressed batch effects using the removeBatchEffect function. Our dataset was analyzed using xCell, including both control and psoriasis samples. A heatmap illustrating the average cell type enrichment scores for each group (PS male, PS female, Control male, Control female) was generated using the ComplexHeatmap package. Boxplots of the enrichment scores for each group were created and compared using the Wilcoxon test, with cell types exhibiting a p-value < 0.05 considered significantly differentially enriched. To assess differences between male and female psoriasis, we calculated Cohen’s D scores to quantify the magnitude of these differences. Cohen’s D measures the effect size, indicating the extent of differences between the psoriasis and control groups for each sex. The scores were calculated by comparing the mean differences in cell type abundance between the psoriasis and control groups, divided by the pooled standard deviation. While Cohen’s D provides insight into the size of these differences, it does not directly assess statistical significance. Statistical significance was determined separately based on differences between disease and control groups. Only cell types that significantly differed between the disease and control groups for each sex were included in the Cohen’s D analysis.

### Statistical Analysis

Each tool used its own statistical methods as described above. For comparisons between the four groups, we applied Kruskal-Wallis analysis followed by Dunn’s test for multiple pairwise comparisons.

## Author contributions

Conceptualization: ES, TZ, MSD; Formal analysis: ES, TL; Investigation: ES, TL, GS, IM, MSD; Methodology: ES, LS, MSD; Supervision: MSD; Writing—original draft: ES; Writing—review & editing: MSD, GS, IM

## Data and Materials Availability

All data necessary to evaluate the conclusions of this study are available for download from the GEO or ArrayExpress repositories using the following accession numbers: GSE117405, GSE121212, GSE171012, GSE176279, GSE183820, GSE205748, GSE249936, GSE41745, GSE47944, GSE54456, GSE67785, GSE83645 and EMTAB6556.

