## Supplementary material for "Sex-Based Transcriptomic Differences in Psoriatic Lesions: A Comprehensive Meta-Analysis"

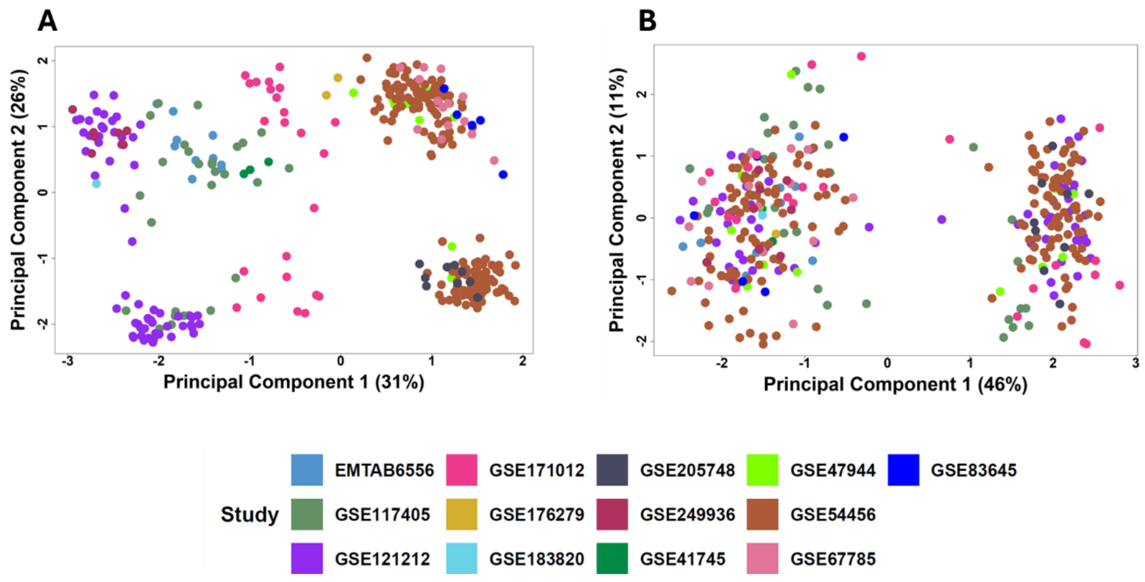


**<u>Supplementary Figure S1:</u>** The Multidimensional Scaling (MDS) plot visualizes the normalized expression of the samples before study batch (A) and after remove study effect(B)

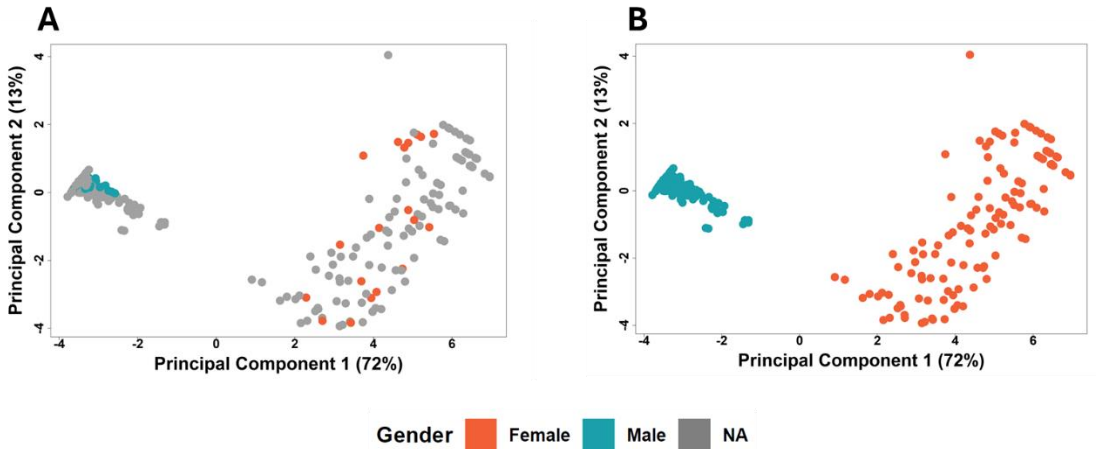


**<u>Supplementary Figure S2:</u>** The Multidimensional Scaling (MDS) plot visualizes the normalized expression of six genes: the female-specific gene XIST and the Y-linked genes EIF1AY, KDM5D, UTY, DDX3Y, and RPS4Y1, to determine sample sex. (A) Before inferred sex (B) After sex inference. Colors represent Male (*Blue*), Female (*Orange*), and NA (*Grey*).

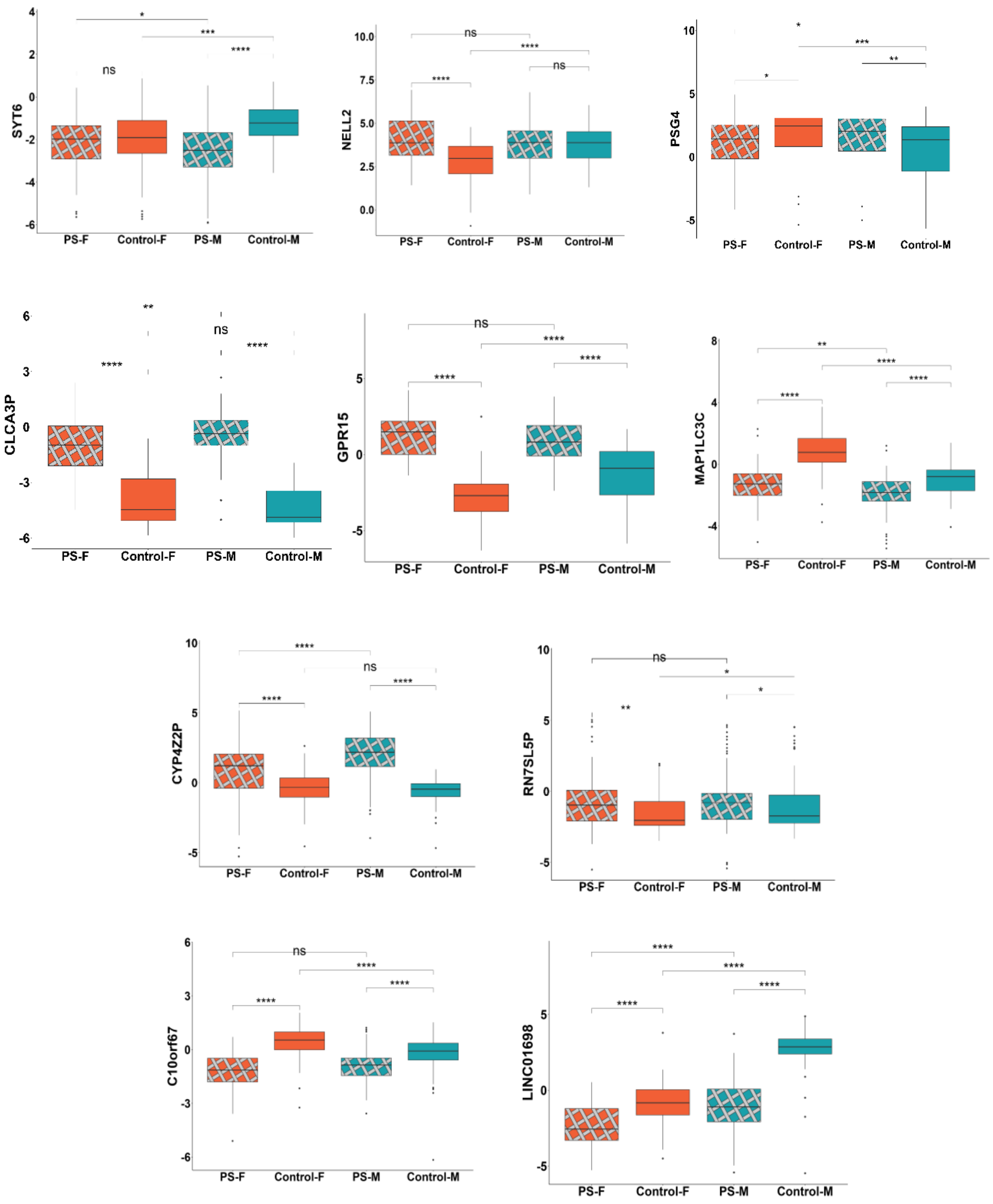


**<u>Supplementary Figure S3</u>:** Boxplot of normalized gene expression after study batch removal, showing the distribution of 10 interaction genes across all cohort groups: Psoriasis female (PS-F), Control female (Control-F), Psoriasis male (PS-M), and Control male (Control-M). Kruskal-Wallis test followed by Dunn's test was used to calculate statistics. Asterisks indicate significance: *$p < 0.05$, **$p < 0.01$, ***$p < 0.001$, ****$p < 0.0001$, and "ns" for non-significant.

A

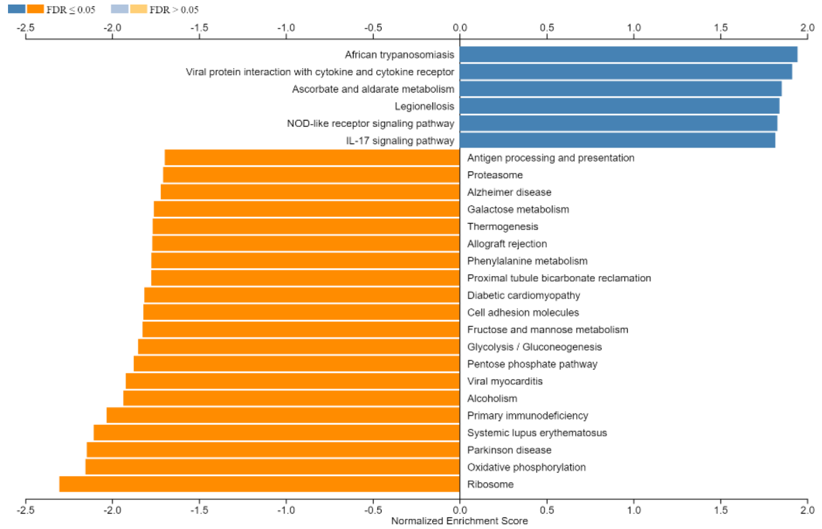


B

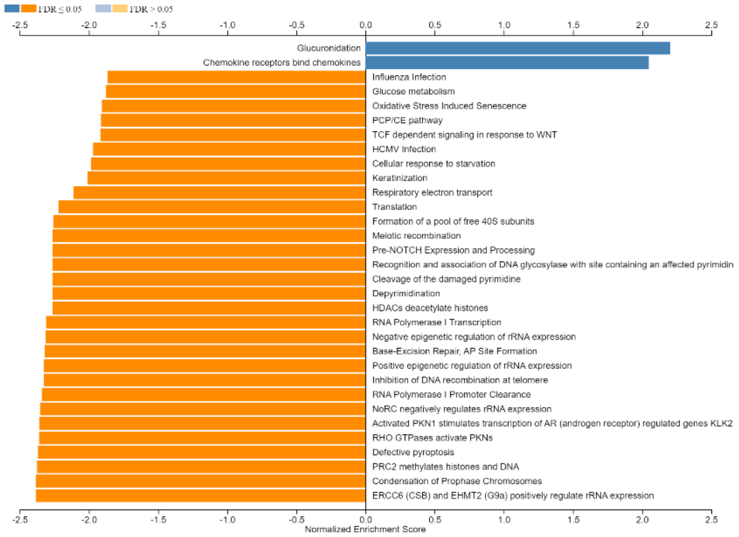


C

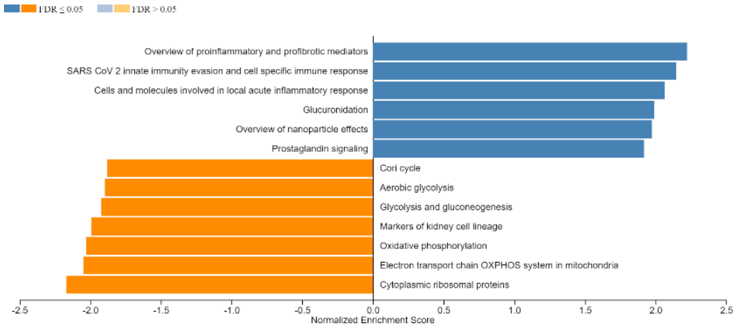


**Supplementary Figure S4**: Barplot showing pathways significantly enriched or depleted in psoriasis males compared to females. Genes were ranked based on the interaction term (PSmale−Controlmale)−(PSfemale−Controlfemale)(PS_male - Control_male) - (PS_female -

Control_female)(PSmale−Control_male)−(PS_female−Control_female), and pathways were identified using GSEA via WebGestalt. Significance was determined using FDR correction (adjusted $p < 0.05$). Bars represent the -log10(FDR-adjusted p-value) of each pathway.Pathway analysis of interaction DEGs using WebGestalt (https://www.webgestalt.org, PMID: 38808672). Results are shown for the KEGG database (A), Reactome database (B), and WikiPathways (C).
